# Cardiac overexpression of microRNA-7 is associated with adverse cardiac remodeling

**DOI:** 10.1101/2020.09.17.302224

**Authors:** Manveen K Gupta, Anita Sahu, Xi Wang, Elizabeth E. Martelli, Kate Stenson, Conner Witherow, Sathyamangla V. Naga Prasad

## Abstract

Role of microRNA-7 (miRNA-7) in targeting Epidermal growth factor receptor (EGFR/ERBB) family is known in dividing cancer cells while less is known about its role in terminally differentiated cardiac cells. We generated transgenic (Tg) mice with cardiomyocyte-specific overexpression of miRNA-7 to determine its role in regulating cardiac function. Despite similar survival, expression of miRNA-7 results in cardiac dilation as measured by echocardiography, instead of age-based cardiac hypertrophy observed in littermate controls. In contrast to the classical adaptive hypertrophy in response to TAC, miRNA-7 Tg mice directly undergo cardiac dilation post-TAC that is associated with increased fibrosis. Interestingly, significant loss in ERBB2 expression was observed in cardiomyocytes with no changes in ERBB1 (EGFR). Gene ontology and cellular component analysis using the cardiac proteomics data showed significant reduction in mitochondrial membrane integrity reflecting the differential enrichment/loss of proteins in miRNA-7 Tg mice compared to littermate controls. Consistently, electron microscopy showed that miRNA-7 Tg hearts had disorganized and rounded mitochondrial morphology indicating mitochondrial dysfunction. These findings show that expression of miRNA-7 uniquely results in cardiac dilation instead of adaptive hypertrophic response to cardiac stress providing insights on adverse remodeling in physiology and pathology.

## Introduction

Despite advancements, symptomatic heart failure is still a chronically progressive disease with therapeutic strategies that only delays outcomes Ref. The heart responds to the increasing demands of the body by undergoing hypertrophic response typically classified as physiological or pathological hypertrophy [14]. Physiological hypertrophy is a reversible response to increased exercise or pregnancy [35], while pathological hypertrophy is irreversible typically leading to cardiac dilation and heart failure [35]. Cardiac hypertrophy is also observed with age and is considered to be a physiological response by the heart to meet the demands of the body. Given that the cardiomyocytes are terminally differentiated, the heart responds to increasing demands of the body by undergoing an adaptive hypertrophic response which over a period of time becomes maladaptive leading to cardiac dilation. Though studies have focused on determining molecular components involved in the transition from hypertrophy to dilation [6, 41] yet, less is known about these pathways. Furthermore, it is known that cancer therapeutics including targeting of epidermal growth factor receptor (anti-ERBB agents) family members [3, 44] results in cardiac dilation of otherwise healthy hearts without an adaptive hypertrophic response [24].

Epidermal growth factor receptors (EGFRs or ERBBs) mediate cardiac hypertrophic response [10] and are represented by four members, ERBB 1 through 4 that either homo- or hetero-dimerize to initiate downstream signals [16]. Studies have shown that ERBB1 (EGFR1) and ERBB2 are transactivated following beta-adrenergic receptor (βAR) activation [36, 46].

While activation of ERBB2 by neuregulin (NRG1) re-initiates cardiomyocyte division [4], conditional ERBB2-knockout mice are characterized by cardiac dilation [11] indicating a key role for ERBB2 in cardiac hypertrophic response. Consistently, cardiomyocyte-specific overexpression of ERBB2 leads to cardiac hypertrophy without heart failure [11, 47]. This further supports the role of ERBB2 in adaptive cardiac hypertrophy as anti-ERBB2 targeted anti-neoplastic therapy in patient’s results in cardiac dilation [13].

Cardiac remodeling encompassing hypertrophic response and dilation is a complex process, in part mediated by microRNAs (miRNAs) [19, 50, 54] miRNAs mediate these effects by binding to complementary mRNA transcripts[2, 25] Despite their ability to bind multiple mRNAs, expression of miRNAs regulate a primary set of proteins resulting in specific phenotypic outcomes. Given that miRNA-7 is known to target the ERBB receptor family in cancer cells [53], and play a role in other disease states [21], we generated a transgenic (Tg) mice with cardiomyocyte-overexpression of miRNA-7 (miRNA-7 Tg) as a strategy to target ERBB in the heart. Consistently, cardiomyocyte expression of miRNA-7 results in significant reduction of ERBB2 that is associated with age-based cardiac dilation and accelerated heart failure post-TAC. An observation, reminiscent of the cardiac dilation observed with anti-ERBB2 agents [9]. However, recognizing that miRNA-7 could have targets beyond ERRB family, proteomics was performed on the miRNA-7 expressing hearts. Gene ontology cellular component networking analysis showed association with mitochondrial dysfunction with relative loss of mitochondrial membrane proteins in miRNA-7 Tg mice. Correspondingly, electron microscopy showed rounded mitochondrial morphology indicating mitochondrial dysfunction. Thus, understanding these pathways will provide insights on cardiac dilation as it is the common exacerbated phenotype of heart failure observed in end-stage human heart patients and in patients undergoing cancer therapeutics [9, 31].

## Methods

### Experimental Animals

All mice used in the studies are in C57Bl/6 background. Transgenic (Tg) mice expressing miRNA-7 and littermate controls were used in the study. <u>Generation of miRNA-7 Tg mice:</u> Mice with cardiomyocyte-specific expression of mature human miR-7-1-5p sequence (5’-UGGAAGACUAGUGAUUUUGUUGU-3’) under alpha-MHC promoter were generated at the Transgenic targeting facility (Case Western Reserve University) ^8^ in the C57BL/6 background. Four positively identified founders were bred with C57BL/6 mice. After back-breeding for three generations, the F3 pups were assessed for expression of miR-7-1-5p in the hearts. Only one founder line showed significant expression of miR-7 which was used in our studies. The Kaplan-Meier procedure was used to calculate survival with time and the data is as % survival over days. Animals were handled according to the approved protocols and animal welfare regulation of IACUC at Cleveland Clinic following the approved NIH guidelines.

### Transverse Aortic Constriction (TAC) and ERBB2 inhibitor studies

miRNA-7 Tg mice and littermate controls were subjected to pressure overload by performing transverse aortic constriction (TAC) surgery. Briefly, following anesthesia with ketamine and xyalzine, the mouse was connected to a rodent ventilator and surgery performed as previously described [36]. Sham operated animals underwent the similar procedure except for the aortic constriction.

### RNA isolation and northern blotting

RNA isolation and northern blotting was performed as described previously [39]. Briefly, left ventricular tissue was homogenized using TRIZOL (Invitrogen) and integrity of the RNA assessed by spectroscopic analysis. 20 μg of RNA was size-fractionated by denaturing formaldehyde gel electrophoresis, transferred to nylon membrane and cross-linked. The membrane was stained with 0.5% methylene blue to visualize the transferred RNA and equal loading. The membranes were de-stained with diethylpyrocarobonate (DEPC)-treated water and hybridized with [^32^P] labeled mature miRNA-7 probe. Following hybridization, the filters were washed under stringent conditions to visualize miRNA-7 transcripts by autoradiography.

### Quantitative real time-PCR

Two μg of total RNA was used for reverse transcription (RT) with RT kit (Applied Biosystems) using specific primer sets supplied by Taqman kit for miRNA-7-1-5p. Quantitative real-time PCR (qPCR) was performed using cDNA with Taqman real time mix in a BioRad iCycler machine. The fold change in the miRNA levels was determined using RNU6B as an internal control. ΔΔCt method was used to compare the miRNA-7 Tg mice with littermate controls. The 2-ΔΔCT method was used as relative quantification strategy for qPCR data analysis [28].

### Echocardiography

Echocardiography was performed on anesthetized mice using a VEVO 770 and VEVO 2100 (VISUALSONICS) echocardiographic machine as previously described [52]. M-mode recording was used to obtain parameters functional parameter.

### Cardiac lysate and Western Immunoblotting

Cardiac lysates were isolated as previously described [51]. Briefly, the cardiac tissue was homogenized in lysis buffer (1% Nonidet P-40 (NP 40), 20 mM Tris-Cl pH 7.4, 300 mM NaCl, 1 mM EDTA, 20% glycerol, 0.1 mM PMSF, 10 μg ml^−1^ each of Leupeptin, and Aprotinin). The cardiac lysates were centrifuged 38,000 × g for 25 min at 4°C and the supernatant used for western analysis. 100 µg of total cardiac lysates were resolved on a SDS-PAGE gel, transferred to PVDF membranes (BIO-RAD) and western immunoblotting was performed using anti-ERBB2 (Neu) antibody (1:500) followed by chemi-luminescence. Similarly, 50 µg of adult cardiomyocyte lysates were used for ERRB2 western immunoblotting studies. Densitometry analysis was carried out using the NIH image J software.

### Isolation of adult cardiomyocytes

The mice were anesthetized; the excised heart was immediately cannulated with 20-gauge needle and mounted to perfusion apparatus. The perfusion buffer contained 113 mM NaCl, 4.7 mM KCl, 0.6 mM KH2PO4, 0.6 mM Na2PO4, 1.2 mM MgSO4, 0.5 mM MgCl2, 10 mM HEPES, 20 mM D-glucose, 30 mM taurine and 20 μM Ca2+ at pH 7.4. Following perfusion for 4 minutes, 150 units/ml of type II collagenase was perfused for 15 minutes. All the solutions were maintained at 34 °C and continuously bubbled with 95% O2 and 5% CO2. Left ventricular tissue was separated from the atria and right ventricle, minced, and digested in perfusate for 15 min. The digested heart was filtered through 200 μm nylon mesh, placed in a conical tube, and spun at 100 rpm to allow viable myocytes to settle. Serial washes were used to remove nonviable myocytes and digestive enzymes, and the adult myocytes were collected. Isolated cardiomyocytes were used for in vitro isoproterenol stimulated cardiomyocyte contractility using the IonOptix System (Myopace, Milton, MA) as previously described [52] and subset were lysed in the lysis buffer for western immunoblotting.

### Immunohistochemistry

Immunohistochemistry was performed as previously described [32]. Excised hearts were fixed in 5% paraformaldehyde for 24 hours, processed for embedding and sectioning at the institutional imaging core. The sections were stained using standard protocol of the imaging core for H & E, Masson’s trichome and Picrosirius Red. <u>Measurement of dilated heart</u> area and area of heart tissue in the transverse sections of the heart were performed using the Image Pro-Plus software at the Cleveland Clinic institutional imaging core. Dilation and heart area were measured in the miRNA-7 Tg and their littermate controls allowing for the Image Pro-Plus software to assess the area of dilation.

### Proteomics,Network analysis and miRNA predicted target analysis

Cardiac lysates from 3 months old miRNA-7 Tg and littermate controls (n=3) were resolved on 10% SDS-PAGE gel and stained with coommassie blue. The stained gels were given to the proteomics core facility within Lerner Research Institute, Cleveland Clinic. In gel digestion was performed and the digested proteins from these sample sets were injected for LC-MS. Relative abundance of the proteins were assessed based on comparing the intensities of the observed peptides. All proteins that have differences in expression of more than two fold were used for the networking analysis. The list of statistically significant unique set of proteins differentially expressed between miRNA-7 Tg versus littermate control were used to query Gene Ontology (GO) - Biological function, Molecular Function and Cellular component database using the ClueGo Cytoscape [5]. The following ClueGo parameters were set wherein, Go Term Fusion would select only display pathways with p values ≤ 0.05; GO tree interval at all levels were set at a term minimum of 3 genes with a threshold of 4% of genes/pathway and a kappa score of 0.42. Gene ontology terms are presented as nodes and clustered together based on the similarity of genes present in each term or pathway. Node size is proportional to the p value for the GO term enrichment. Proteins are presented as circles wherein, red circles indicate upregulated proteins and blue circles indicate downregulated proteins associated with one of more processes. Predicted targets of miRNA-7 were identified using the publicly available miRNA prediction tool miRDB [8] and TargetScan [1]. Analysis of predicted targets for miRNA-7 using miRDB and TargetScan tools showed 758 and 508 targets respectively. These predicted target proteins were compared with our proteomics data showing differential expression of proteins in miRNA-7 Tg versus littermate controls. A Venn diagram was constructed to show targets that are common between predicted versus experimentally identified target proteins. These set of common proteins present in the proteomics data and prediction tools are represented in the Table 3.

### Transmission Electron Microscopy

Transmission Electron microscopy on heart tissue was performed at the Institutional electron microscopy core. Heart sample was fixed in 2.5% glutaraldehyde/4% paraformaldehyde in 0.2 M sodium cacodylate buffer overnight at 4° C.

Sample was washed three times, five minutes each with 0.2 M sodium cacodylate buffer (pH 7.3). After wash with the buffer, cold water dissolved 1% Osmium Tetroxide was added, followed by sodium cacodylate buffer wash and rinsed with Maleate buffer (pH 5.1). The tissue was stained with 1% uranyl acetate in Maleate buffer for 60 minutes, washed with maleate buffer and dehydrated by washing with increasing concentrations of cold ethanol from 30% through 95% once each time for 5 minutes followed by 100% ethanol three times for 10 minutes each at room temperature. The tissue was incubated in propylene oxide for 15 minutes each for three times and propylene oxide was removed by overnight incubation with 1:1 propylene oxide/eponate 12 at room temperature followed by pure eponate 12 medium for 4-6 hours. The tissue sample was polymerized and semi-thin section of 1 µM were cut with diamond knife stained with toluidine blue for observation in a Leica DM5500 light microscope. Ultra-thin sections of 85 nm were cut with diamond knife, stained with uranyl acetate and lead citrate, and observed with a Tecnai G2 SpiritBT, electron microscope operated at 60 kV.

### Statistics

Data are expressed as mean + SEM. Statistical comparisons were performed using an unpaired Student’s *t*-test for two samples comparison and analysis of variance (ANOVA) was carried out for multiple comparisons for paired echocardiography parameter analysis. Post-hoc analysis was performed with a Scheffe’s test. For analysis, a value of * p< 0.05 was considered significant.

## RESULTS

### Cardiomyocyte-specific overexpression of miRNA-7 leads to cardiac dilation

To test whether miRNA-7 Tg mice express miRNA-7, northern blotting was performed on RNA from the F3 pups miRNA-7 Tg and littermate controls. Significant expression of miRNA-7 in the miRNA-7 Tg mice was observed compared to littermate controls (NTg) [**Fig. 1a**]. To further confirm the expression of miRNA-7, quantitative real time PCR (qRT-PCR) was performed that showed significant expression of miRNA-7 in Tg mice [**Fig. 1b**]. Since miRNA-7 Tg mice displayed no overt abnormality in life span, H & E staining was performed on heart sections of 12 month old miRNA-7 Tg and NTg mice. Sections from miRNA-7 Tg mice show marked dilation compared to NTg [**Fig 1c**]. Measurement of heart tissue area using Image Pro-Plus showed significant dilation in the miRNA-7 Tg mice [**Fig. 1d**] and was inversely correlated to decreased cardiac tissue area compared to age-matched NTg (12 months) (see methods) [**Fig. 1e**]. As cardiac dilation is observed in 12 month miRNA-7 Tg mice, survival analysis was performed over 20 month period. Interestingly, Kaplan-Meier survival curves show no appreciable differences in survival rates between miRNA-7 Tg and NTg mice [**Fig. 1f**].

**Figure 1.**
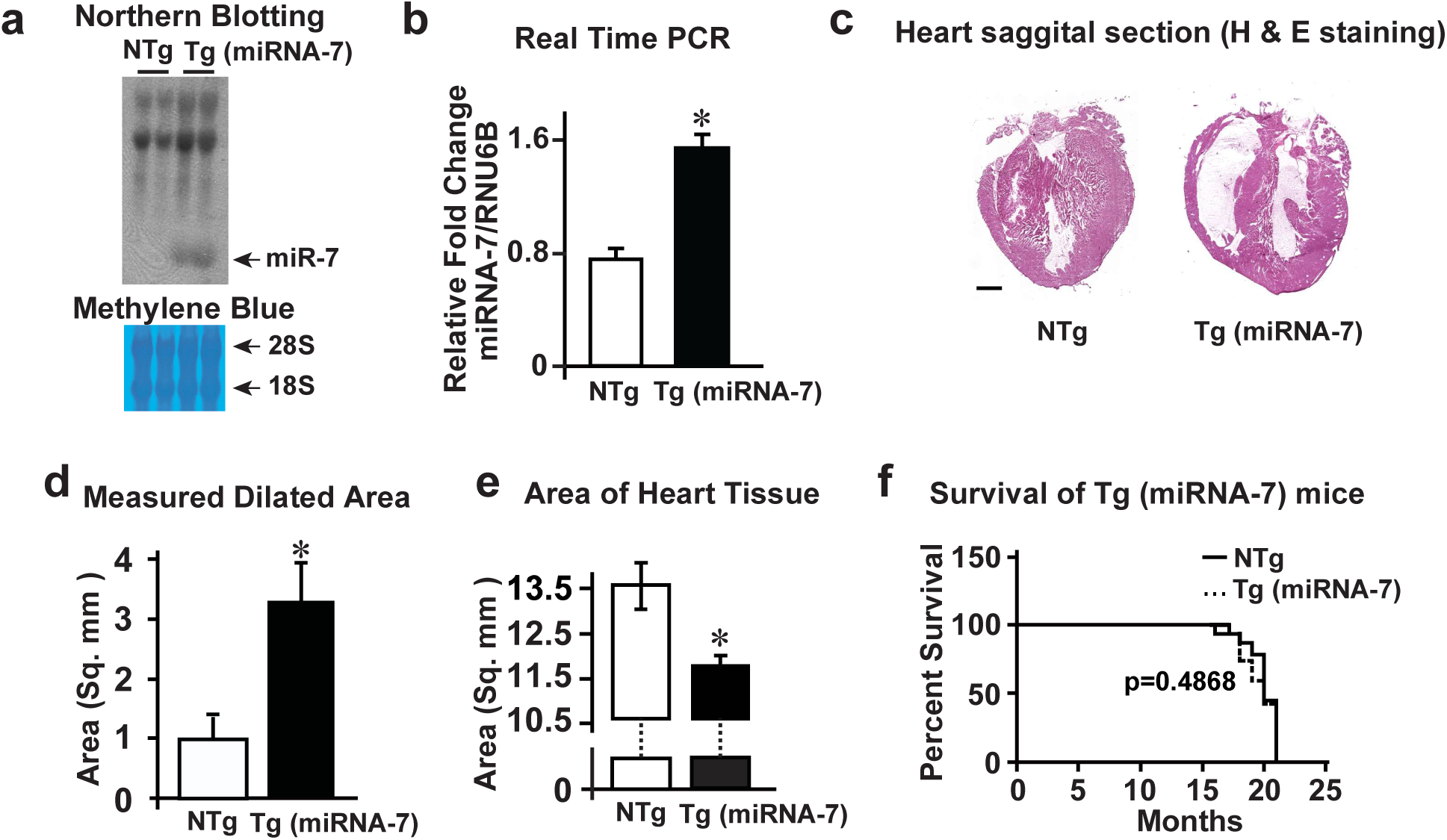
microRNA-7 (miRNA-7) Tg mice have similar life span despite cardiac dilation. **a**, Northern blotting for miRNA-7 in the Tg (miRNA-7) mice and littermate controls (NTg) (upper panel); Methylene blue staining of RNA for equal loading (lower panel) (n=5). **b**, Real time PCR analysis for miRNA-7 expression in the NTg and Tg mice. *p<001 vs. NTg (n= 5). **c**, Cardiac sections from 12 month old NTg and Tg mice stained with H & E wherein, Tg hearts are characterized by marked dilation compared to NTg (n=4). Scale bar 1000 µm. **d & e**, The dilated area (d) and the area of cardiac tissue (e) was measured with Image Pro Plus using the 12 month old H & E stained heart sections (n=4). *p< 001 vs. NTg (d) & *p< 0.01 vs. NTg (e). **e**, Kaplan-Meier survival curves for NTg and Tg mice shows no appreciable differences in survival between NTg and Tg (n=18).

### Increasing cardiac dysfunction with age in miRNA-7 Tg mice

Given the similar survival rates despite cardiac dilation in miRNA-7 Tg mice, we investigated the cardiac function by echocardiography in miRNA-7 Tg and NTg over 18 months. M-mode echocardiography and functional parameters (left ventricular end-systolic dimeter, LVESD; LV end-diastolic diameter, LVESD; % fractional shortening, % FS) showed that miRNA-7 Tg mice has significant cardiac dysfunction by 12 months of age that deteriorates further by 18 months [**Fig. 2a**,**b and Table
 1**]. Morphometry also showed significant increase in HW/BW ratios in miRNA7-Tg compared to NTg mice [**Table 1**] reflecting adverse cardiac remodeling and exacerbation of cardiac dysfunction due to dilation in miRNA-7 Tg mice.

**Figure 2.**
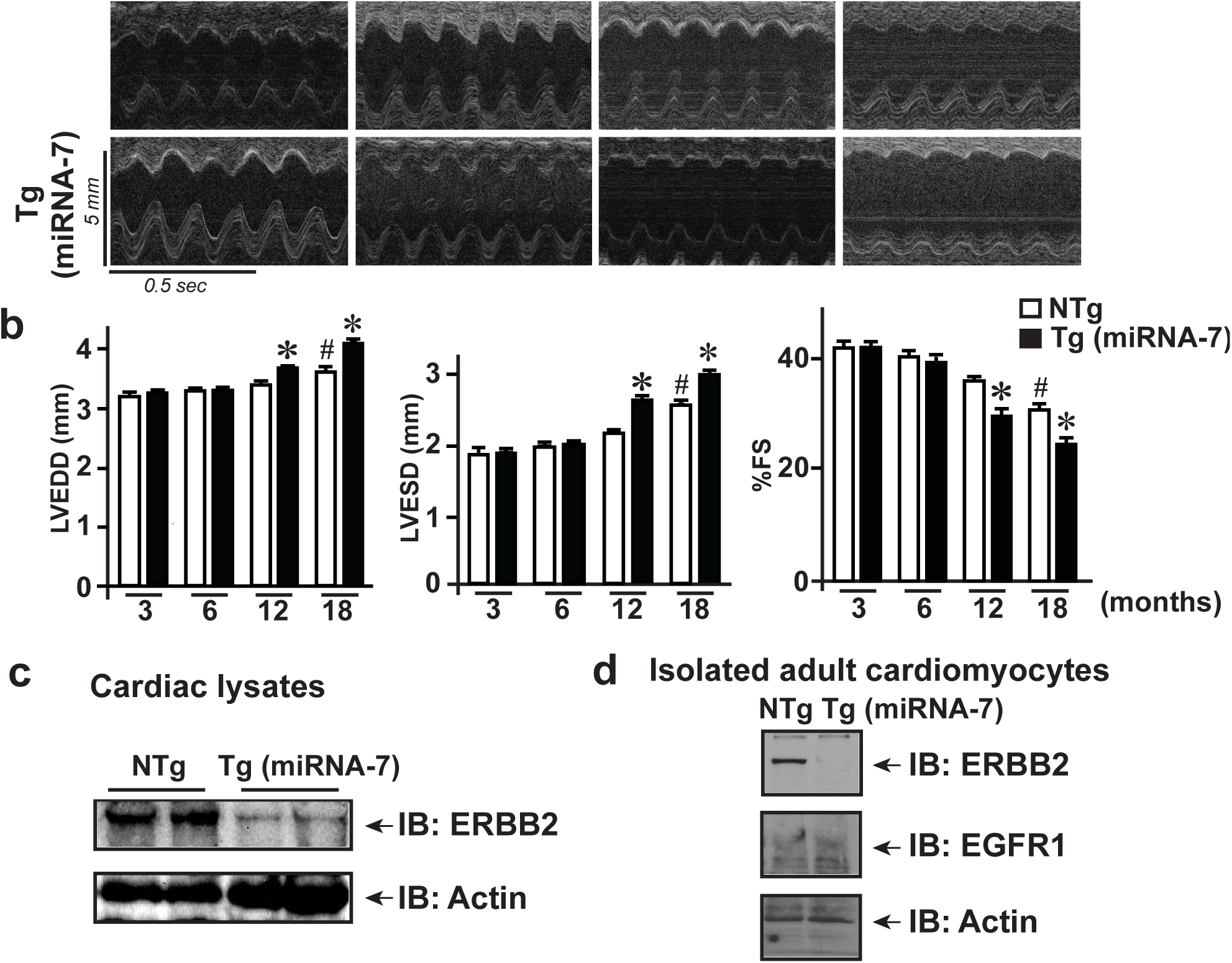
Cardiomyocyte expression of miRNA-7 leads to age-based cardiac dilation. **a**, M-mode echocardiography performed on NTg and Tg mice at 3, 6, 12 and 18 months (n=12). **b**, Cardiac functional parameters measured from echocardiography on NTg and Tg mice; Left panel: LVEDD (left ventricular end-diastolic dimension); Middle panel LVESD (left ventricular end-systolic dimension); Right panel: % fractional shortening (%FS) as measure of cardiac function. *p<0.01 vs. 3 month NTg and Tg; # p<0.05 vs. 3 month NTg and Tg. **c**, Immunoblotting for ERBB2 in the total cardiac lysates from NTg and Tg mice (n=8). **d**, Immunoblotting for ERBB2 and EGFR in isolated adult cardiomyocytes from NTg and Tg mice (n=5).

### miRNA-7 Tg hearts have reduced ERBB2 levels

Studies have shown the members of the epidermal growth factor receptor (EGFR) family play a key role in cardiac hypertrophy and growth response [38, 47]. As miRNA-7 targets ERBB receptor [53], total cardiac lysates were immunoblotted for ERBB2. Marked reduction in ERBB2 expression was observed in miRNA-7 Tg mice compared to NTg [**Fig. 2c**]. To provide unequivocal evidence that miRNA-7 expression targets ERBB2 in the cardiomyocytes, adult cardiomyocytes were isolated form miRNA-7 Tg and NTg hearts. Cardiomyocyte lysates were immunoblotted for ERBB2 and EGFR (ERBB1). Consistently, there was significant reduction in ERBB2 expression in miRNA-7 Tg cardiomyocytes compared NTg [**Fig. 2 d**]. While, we did not observe any appreciable differences in EGFR (ERBB1) expression between miRNA-7 Tg and NTg mice [**Fig. 2 d**] suggesting that the adverse cardiac remodeling observed with age in miRNA-7 Tg mice could in part, be due to reduced expression of ERBB2.

### Pressure overload hypertrophy by transverse aortic constriction (TAC) accelerates cardiac dysfunction in miRNA-7 Tg mice

Since the miRNA-7 Tg mice survive normally despite cardiac dysfunction, we assessed whether subjecting older mice (12 months) to pressure overload by TAC would accelerate their cardiac dysfunction/survival. Following TAC, all miRNA-7 Tg mice died within 4 days while no deaths were observed with NTg [**Fig. 3a**] showing that older miRNA-7 Tg cannot withstand pressure overload induced by TAC. To test whether younger mice would with stand the pressure overload by TAC, 3 months miRNA-7 Tg and NTg mice were subjected to TAC for two weeks. Although the 3 month old miRNA-7 Tg mice survived the TAC, they underwent severe adverse remodeling that was associated with significant cardiac dilation and dysfunction. Gravimetric analysis showed significant increase in heart weight to body weight (HW/BW) ratio in miRNA-7 Tg mice compared to NTg [**Fig. 3b**]. While no appreciable differences were observed in baseline echocardiography measurements, subjecting miRNA-7 Tg to TAC significantly accelerated cardiac dysfunction resulting in cardiac dilation [**Table 2**]. Echocardiography assessment showed that miRNA-7 Tg mice were characterized by cardiac dilation associated with increased left ventricular end-diastolic and -systolic dimensions (LVEDD and LVESD) post-TAC [**Fig. 3 c & d**]. Consistent with cardiac dysfunction, miRNA-7 Tg mice displayed significant reduction in % factional shortening (%FS) and % ejection fraction (%EF) post-TAC compared to NTg [**Fig. 3 e & f**]. In contrast to the cardiac dilation observed in the miRNA-7 Tg, NTg mice showed classical adaptive hypertrophic response to TAC as reflected by increased anterior and posterior wall thickness [**Table 2**]. Corresponding to cardiac dilation, miRNA-7 Tg mice have decreased anterior and posterior wall thickness [**Table 2**]. These observations show that miRNA-7 expression exacerbates cardiac dilation in part, through ERBB2 regulation indicating. However, given that miRNA-7 could target other proteins, the overall phenotype could be a determined by contributions of other proteins altered by miRNA-7 in addition to ERBB2. Given the observation of cardiac dilation in response to pressure overload, in vitro adult cardiomyocyte contraction in the miRNA-7 Tg and NTg were performed to assess for dysfunction. Adult cardiomyocytes were isolated from 3 month miRNA-7 Tg and NTg mice and in vitro contractility assessed in response to isoproterenol (ISO). Cardiomyocytes from NTg mice had a classical response to ISO [**Supplementary Fig. 1a, upper right panel (ISO) & 1b**]. Although the baseline myocyte contractility in miRNA-7 Tg mice is similar to NTg [**Supplementary Fig. 1a, lower left panel (baseline)**], there was a significant reduction in ISO response [**Supplementary Fig. 1a, lower right panel (ISO) & 1b**]. These data further supports the idea that miRNA-7 Tg mice have an innate predisposition to cardiac dysfunction which is exacerbated in response to stress.

**Figure 3.**
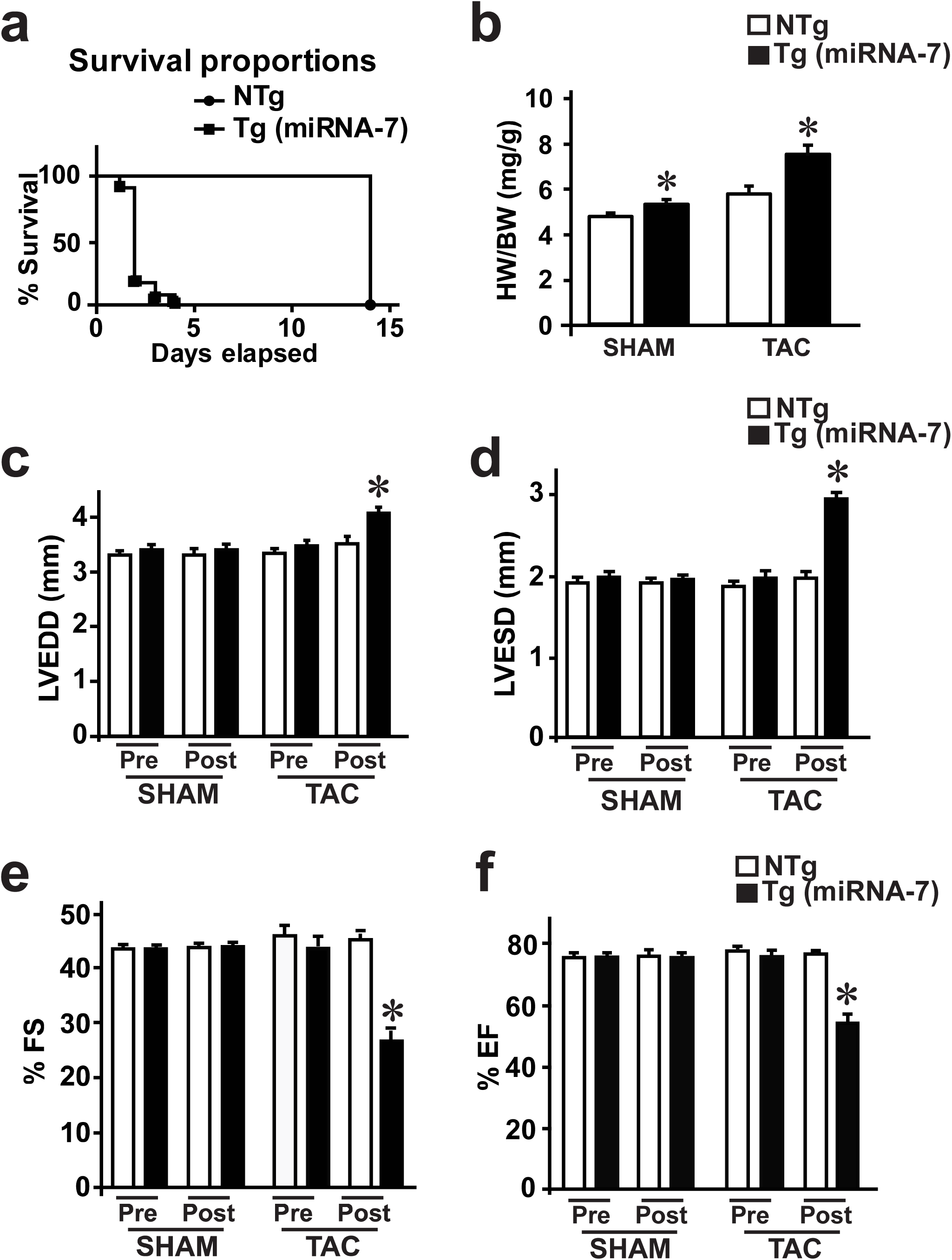
Cardiomyocyte expression of miRNA-7exacerbates cardiac dysfunction and failure following pressure overload by transverse aortic constriction (TAC) **a**, Kaplan-Meier survival curves for 12 months old NTg and Tg mice subjected to TAC (n=8). **b**, Heart weight to body weight (HW/BW) ratio of NTg and Tg following two weeks of TAC on younger mice (3 months) (n=11). **c-f**, Cardiac functional parameters of NTg and Tg mice measured by echocardiography following 2 weeks of TAC (n=11); LVEDD (left ventricular end-diastolic dimension) (**c**), LVESD (left ventricular end-systolic dimension) (**d**), percent fractional shortening (%FS) (**e**), percent ejection fraction (%EF) (**f**). *p<0.005 vs. Sham (NTg or TG) and pre-TAC (NTg or Tg) (n=11).

### Dilation in miRNA-7 Tg mice is associated with fibrosis

Because miRNA-7 Tg mice undergoes cardiac dilation instead of the adaptive hypertrophy in response to TAC, immunohistochemistry (H & E, Masson’s trichrome and picrosirus red staining) was performed to determine whether accelerated cardiac dilation is associated with fibrosis. Consistent with increase HW/BW ratio, miRNA-7 Tg sham heart was slightly larger than NTg sham [**Fig. 4a, panels 1 & 3**]. NTg heart showed the classical adaptive cardiac hypertrophy following TAC [**Fig. 4a, panel 2**]. miRNA-7 Tg had marked increase in size associated with larger lumen [**Fig. 4a, panel 4**] following TAC reflecting cardiac dilation consistent with echocardiography analysis [**Fig. 3**]. Higher magnification H & E staining shows an increase in the cardiomyocyte size in the NTg following TAC [**Fig. 4b, panel 2**] while there are marked disorganization of the cardiomyocytes even in sham miRNA-7 Tg [**Fig. 4b, panel 3 & 4**]. miRNA-7 Tg has elevated Masson’s Trichrome and picrosirius red staining in both sham and TAC hearts [**Fig. 4b, panels 7, 8, 11 and 12**] compared to NTg [**Fig. 4b, panels 5**,**6**,**9 and 10**] showing increased fibrosis. These observations show elevated deposition of interstitial collagen underlying cardiac fibrosis reflecting the exacerbated cardiac dysfunction in miRNA-7 Tg mice [40].

**Figure 4.**
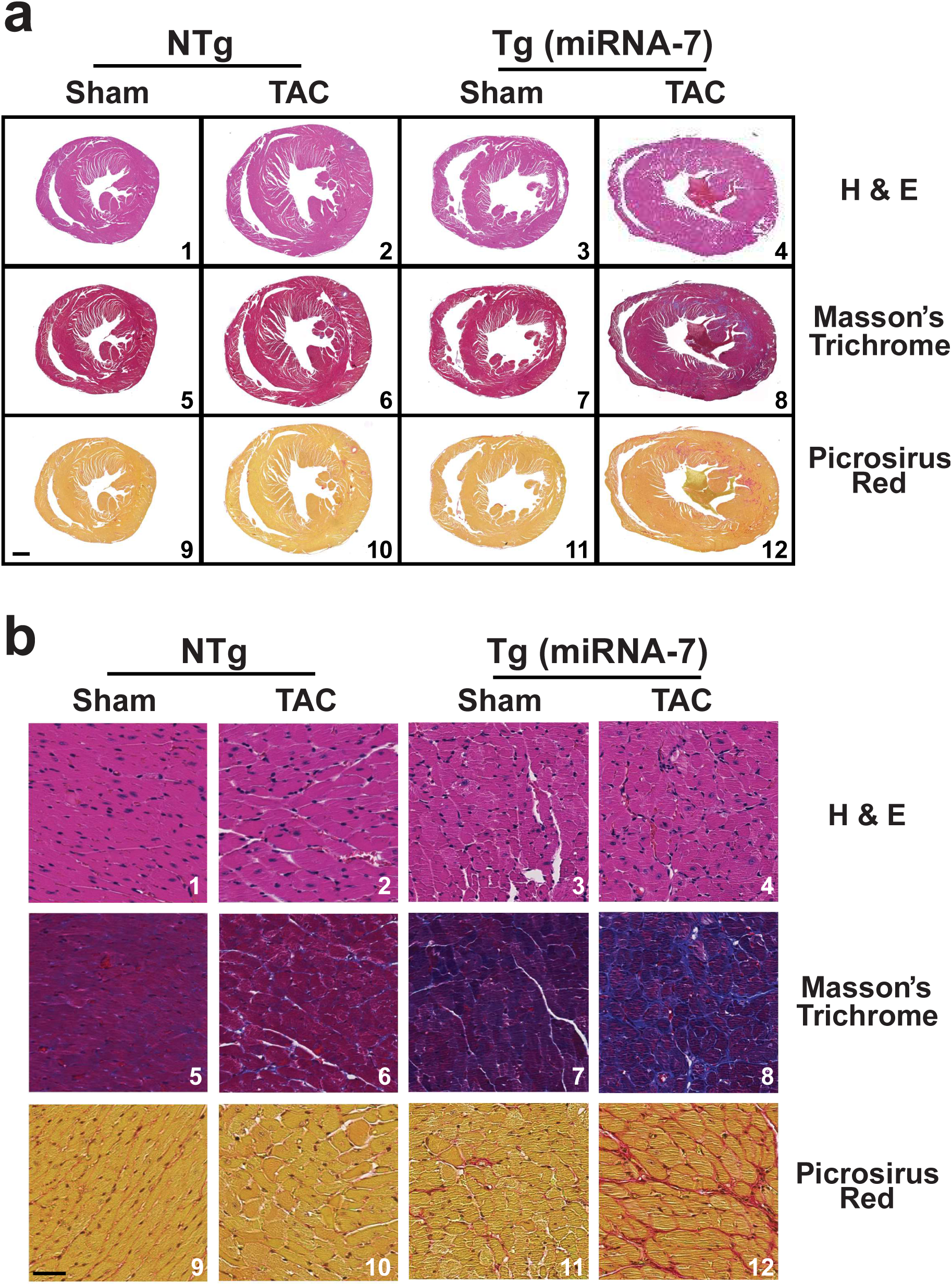
Immunohistochemistry on cardiac sections to assess fibrosis. Heart sections from NTg and Tg were stained with H & E, and collagen deposition was assessed using Masson’s Trichrome or Picrosirius red as indicators of fibrosis (n=5). **a**, Upper panel - lower magnification of the transverse sections. Scale bar 1000 µm. **b**, Lower panel - higher magnification. Scale bar 200 µm (n=5).

### Proteomic and networking analysis in miRNA-7 Tg hearts shows alterations in mitochondrial components

Although we observed loss of ERBB2 expression in the cardiomyocyte following miRNA-7 expression, it is well known that miRNAs can target multiple transcripts and the overall phenotype is a net representation of these effects. In recognition, we performed proteomics on 3 month old hearts from miRNA-7 Tg and NTg mice (n=3) with the rationale that molecular changes precede the phenotypic effects given that cardiac function is still preserved in miRNA-7 Tg mice and similar to NTg [**Fig. 2a**]. Cardiac lysates were resolved on SDS-PAGE, in-gel trypsinized and subjected to LC-MS mass spectrometry analysis. Analysis showed that > 249 proteins were significantly altered with majority of them being downregulated in miRNA-7 Tg hearts compared to NTg despite preserved cardiac function. These set of altered proteins in miRNA-7 Tg hearts were used for gene ontology (GO) networking analysis (see methods) to assess for their role in molecular function and cellular component enrichment. Molecular functional analysis of the altered proteins in miRNA-7 Tg mice showed upregulation of pathway involved in phagocytosis and actin remodeling while there are significant downregulation of oxidative phosphorylation associated with loss in NADH dehydrogenase complex assembly [**Fig. 5a**]. To further dissect the changes in molecular function, cellular component analysis was performed. Examination of GO cellular components showed marked upregulation of proteasomal pathways and significant downregulation mitochondrial matrix and envelope components in miRNA-7 Tg hearts compared to NTg [**Fig. 5b]**. To directly assess how many of these altered proteins in proteomics represent direct targets of miRNA-7, we used two miRNA target predicting databases (miRDB and TargetScan) to compare the predicted targets versus the actual representation of proteins downregulated in hearts. The predicted and the proteomics data is represented in the Venn diagram showing that among > 249 altered proteins, there were only 18 proteins that represented the predicted targets of miRNA-7 and were significantly downregulated in miRNA-7 Tg hearts compared to NTg. Interestingly, among these 18 only 7 downregulated proteins [**Table 3**] were common to both the miRDB and TargetScan prediction databases [**Fig. 5c**]. Among the 7 downregulated proteins, one of the targets GHITM (Growth hormone-inducible transmembrane protein) is interesting it seems to play a role in cristae organization, and its downregulation results in mitochondrial fragmentation [27]. GHITM targeting by miRNA-7 in part, may also contribute to cardiac dysfunction in the Tg mice and would be consistent with the GO cellular component analysis showing significant downregulation of mitochondrial matrix and envelope components that may underlie mitochondrial dysfunction.

**Figure 5.**
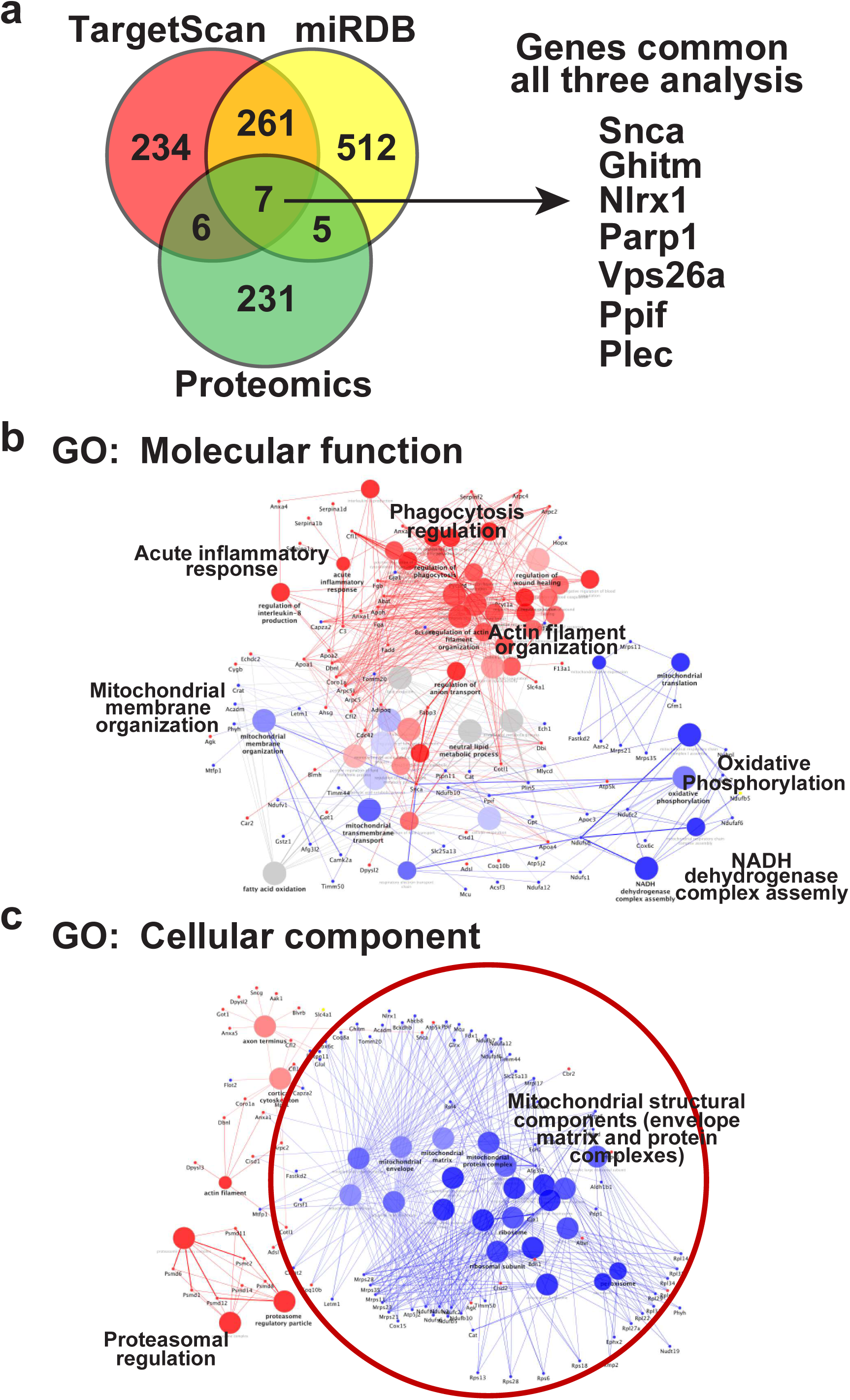
Cardiac proteomics and network analysis of miRNA-7 targets in NTg and Tg hearts. **a**, Venn diagram showing an overlap of predicted targets (from prediction databases TargetScan and miRDB) and truly altered proteins identified in the cardiac proteomic studies from NTg and Tg hearts. Seven common proteins identified are shown the right panel and all are downregulated in the miRNA-7 Tg mice compared to Tg. **b**, Gene ontology (GO) functional analysis using the significantly altered proteins in the Tg versus NTg from cardiac proteomics shows unique molecular functional changes in the NTg wherein phagocytosis, actin remodeling and acute inflammation are upregulated (red color coding). While, mitochondrial structure and function are downregulated (blue color coding). **c**, GO cellular component analysis shows upregulation of proteosomal components (red color) while, majority of the pathways are associated with downregulation of mitochondrial structural components.

### Altered mitochondrial ultrastructure in miRNA-7 Tg hearts

To test whether miRNA-7 Tg mice have mitochondrial structural abnormalities that in turn may underlie its dysfunction, transmission electron microscopy was performed on the 3 months old miRNA-7 Tg and NTg hearts. Significant difference in structure and morphology of the mitochondria was observed in the miRNA-7 Tg hearts compared to NTg [**Fig. 6**]. Mitochondria are significantly disorganized in the miRNA-7 Tg hearts wherein the mitochondria are round with ultrastructure showing lack of cristae that seem to be undergoing fission. Together these observation suggest changes in mitochondrial structure may contribute to mitochondrial dysfunction and in part may leads adverse remodeling.

**Figure 6.**
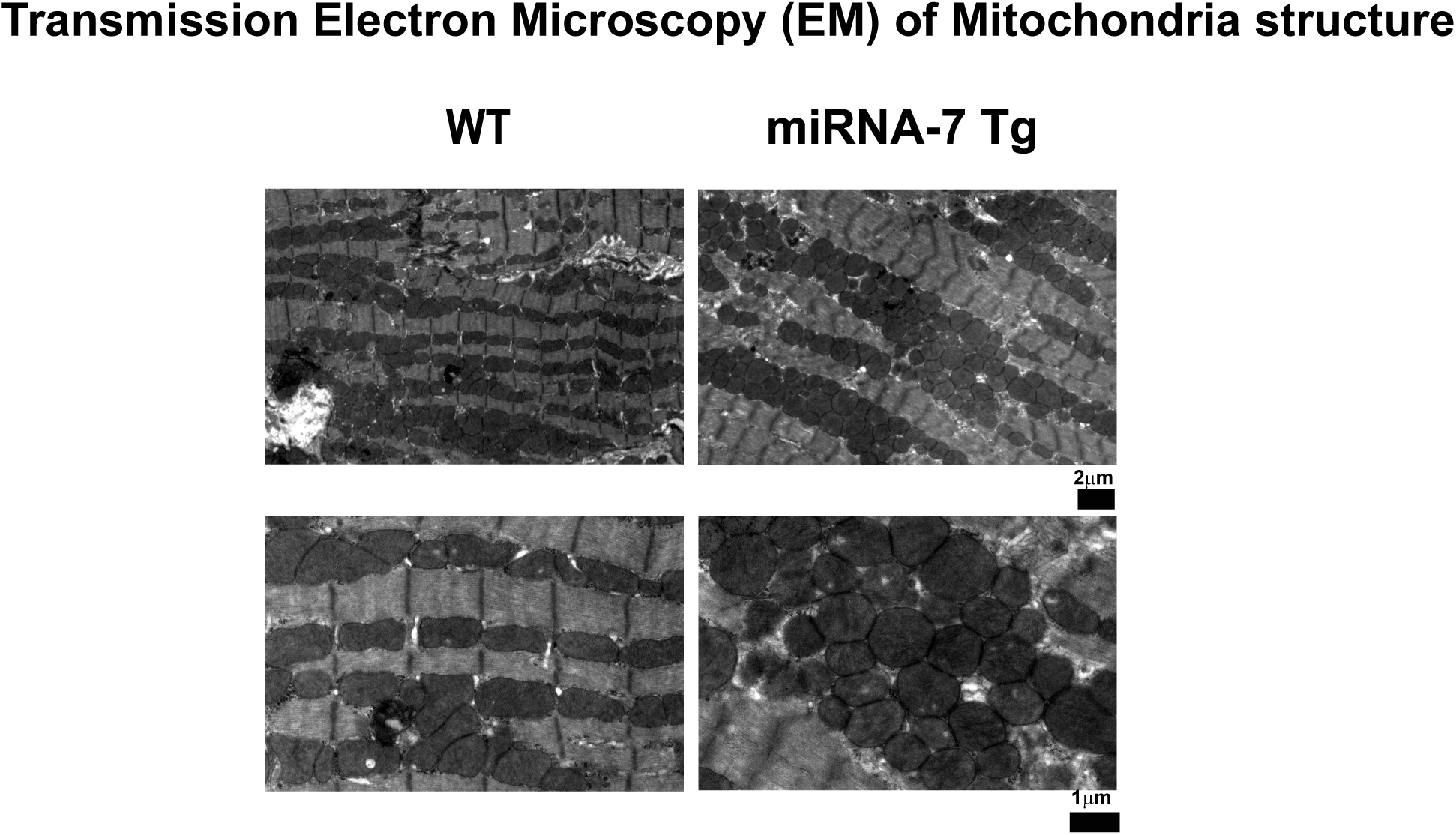
microRNA-7 (miRNA-7) Tg mice are characterized by structurally altered mitochondria. Transmission electron microscopy of the heart sections from NTg and Tg shows marked alteration in mitochondrial ultra-structure. NTg mic are characterized by well-organized and mitochondria aligned with sarcomeres. Tg mitochondria are disorganized and rounded in shape reflecting alterations in mitochondrial structural components (n=5). Upper panel - lower magnification (scale bar - 1μm); Lower panel - higher magnification (scale bar - 2 μm).

## DISCUSSION

Traditionally, heart undergoes adaptive hypertrophic response to age (considered physiological) and to pressure overload (pathological) [49]. However, our study shows that cardiomyocyte expression of miRNA-7 leads to cardiac dilation by-passing the classical adaptive hypertrophy to mechanical overload. This is supported by the findings that miRNA-7 Tg mice undergo age-based cardiac dilation in contrast to NTg characterized by physiological adaptive hypertrophy. The younger miRNA-7 Tg mice despite having similar cardiac function like NTg, directly transition to dilation following TAC indicating a unique role for miRNA-7 in this process. More importantly, cardiac dilation in miRNA-7 Tg mice post-TAC is associated with increased fibrosis as assessed by Masson trichrome staining and collagen deposition by picrosirus red. Intriguingly, miRNA-7 expression leads to significant loss in ERBB2 expression in the cardiomyocytes instead of the EGFR that is observed in cancer cells. Furthermore, proteomic analysis performed on the miRNA-7 Tg hearts surprisingly, showed only 7 miRNA predicted target proteins to be significantly downregulated including protein that play a role in mitochondrial integrity. Consistent with this observation, high resolution electron microscopy showed rounded and disorganized mitochondria that in part, could also play a role in exacerbated cardiac dysfunction observed in miRNA-7 Tg mice.

Cardiomyocyte-specific miRNA-7 expression results in abrogation of ERBB2 expression in cardiomyocytes underlying cardiac dilation, a phenotype consistent with the conditional ERBB2 knockout mice [11, 15]. Despite significant ERBB2 abrogation in cardiomyocytes, total cardiac lysates show low levels of ERBB2 expression reflecting contributions from other cell types in the heart. Interestingly, studies in cancer cells have shown that miRNA-7 reduces EGFR (ERBB1) expression [53]. While miRNA-7 expression in the cardiomyocytes does not alter EGFR expression but abrogates ERBB2 expression reflecting on cell-specific regulation of ERBB1 and ERBB2 despite containing the miRNA-7 targeting leader/seed sequence. This unique regulation could be due to the repertoire of cell-specific mRNA binding proteins that would mask the miRNA binding site on mRNA [17, 18]. This critical component has been overlooked in miRNA studies as expression of different RNA binding proteins in a given cell could contextually change the effects of miRNA on target protein expression [18].

It is interesting to note that miRNA-7 Tg mice display normal cardiac function at younger age but transitions to dilation with age instead of the adaptive cardiac hypertrophy observed in littermate controls with age. Age-based hypertrophic response is considered physiological [7] but however, it is not reversible [14]. This response plays a key role in the initial adaptive hypertrophy to mechanical demands [35] and our observation suggests that this response in part, could be mediated by ERBB2 as loss in ERBB2 expression in miRNA-7 Tg mice leads to cardiac dilation. In this context, chemotherapeutics including anti-ERBB2 agents in human leads to cardiac dilation [9, 45] of the otherwise healthy hearts suggesting a role for ERBB2 pathways in maintaining homeostatic cardiac hypertrophic response. Moreover, ERBB2 expression also seem to in part contribute towards mediating post-TAC adaptive hypertrophic response as its absence in the miRNA-7 Tg mice results in cardiac dilation and accelerated cardiac dysfunction. Such a role in supported by the observation of cardiac dilation in young conditional ERBB2 knockout mice [38] and critically the observation of cardiac hypertrophy with no failure in transgenic mice with cardiomyocyte-specific overexpression of ERBB2 [47]. Thus, in addition to its role in embryonic development [16], cardiomyocyte de-differentiation and proliferation [12], these observations in mouse model elucidates an understudied unique role for ERBB2 in adaptive hypertrophic response that is homeostatic in nature.

miRNA-7 expression consistently targets ERBB2 but however it has to be recognized that miRNA-7 can target other mRNA transcripts to reduce their protein expression [21, 34] which together may contribute to adverse cardiac phenotype. In recognition, global proteomics was performed to assess altered proteins in miRNA-7 Tg mice and NTg littermate controls. Although > 249 proteins were significantly altered, only 18 of the predicated targets (using miRDB and TargetScan) were present in our cardiac proteomics [**Table 3**]. Among these 18 proteins only 7 were common between miRDB and TargetScan prediction database and consistently, all them as are downregulated [**Table 3**]. Analysis of these target proteins show that except for poly(ADP-ribose)polymerase-1 (PARP1) whose role is known in cardiovascular disease [20], nothing is known about the roles of other proteins in cardiac function and regulation. Given that PARP1 inhibition is beneficial in ischemia-reperfusion and our current observation that PARP1 is downregulated in miRNA-7 TG mice and yet these mice have exacerbated heart failure indicating that PARP1 may not a determinant in the overall deleterious phenotype. Similarly, SNCA (alpha-synuclein) accumulation is thought to underlie sympathetic denervation that is associated in patients with Parkinson Disease [22]. While its accumulation causes pathology, it downregulation in miRNA-7 Tg may not have effects on adverse cardiac remodeling. Similarly, our proteomics networking functional analysis showed marked increase in actin cytoskeleton and intracellular trafficking components that could be a reflection of compensatory effects on downregulation of VPS26a that plays a key role in endosomal cargo sorting [23, 29]. Plectin gene (Plec1) polymorphism is known to be associated to hypertrophic cardiomyopathy in humans [48] but its role in cardiomyocytes still needs to be determined. Plec1 mutations are known to underlie muscular dystrophy due to altered interactions with mutant PLEC1 protein to cytoskeleton [42]. However, despite the loss of PLEC1 protein in miRNA-7 Tg mice, there was no disorganization of the sarcomeres and the mice underwent cardiac dilation instead of hypertrophic myopathy observed Plec1 mutations [48].

Critically, three of the downregulated proteins in miRNA-7 Tg hearts are localized to mitochondria (GHITM, NLRX1 and PPIF). PPIF codes for cyclophin D whose inhibition is considered to beneficial [26] and PPIF is already downregulated in miRNA-7Tg mice indicating that despite its loss these mice undergo adverse remodeling. However, less is known about the roles of NLRX1 or GHITM. NLRX1 is mitochondrial targeted NOD-like receptor family member that plays a role in anti-viral immunity [33] by binding to mitochondrial antiviral signaling protein (MAVS) [43]. Consistent with its loss, our molecular functional analysis shows upregulation of acute inflammatory response indicating a potential role for NLRX1 in the miRNA-7 Tg hearts. Perhaps the most intriguing protein in our proteomics study is the GHITM (Growth hormone-inducible transmembrane protein) also known as transmembrane BAX inhibitor motif containing protein 5 (TMBIM5) [27]. GHITM is a member of the BAX inhibitor containing (TMBIM) family that localizes to the inner mitochondrial membrane (IMM) where it plays a role in apoptosis by mediating alterations in mitochondrial morphology and cytochrome c release [27].

Although the role for GMITM is not known in cardiac systems, it is believed that GHITM maintains cristae organization wherein its downregulation results in mitochondrial fragmentation potentially through fusing of the cristae structures. Thus, the observation of rounded and disorganized mitochondria in miRNA-7 Tg mice suggests a key role for GHITM in maintaining mitochondrial integrity. Furthermore, it is also known that loss in GHITM expression leads to release of pro-apoptotic protein cytochrome C [37]. Thus, the release of cytochrome C from the mitochondria of miRNA-7 Tg mice may result in increased apoptosis that may underlie the fibrosis observed in these mice hearts even at baseline that is exacerbated following TAC [**Fig. 4b**]. Furthermore, GHITM is thought to be responsible for cross-linking cytochrome C to IMM thereby, delaying the release of cytochrome C independent of the permeabilization of the outer mitochondrial membrane. Thus, our proteomic study suggests a role for GHTIM in maintaining cardiac mitochondrial integrity and its loss in miRNA-7 Tg mice leads to rounded disorganized mitochondria and in addition, its role in apoptosis may in part underlie the fibrotic phenotype observed in miRNA-7 Tg mice.

Our comprehensive study shows a miRNA-7 expression leads to adverse cardiac dilation instead of adaptive hypertrophy to stress that in part could be mediated by loss in ERBB2 and GHITM expression. The observation of ERBB2 targeting by miRNA-7 instead of EGFR1 observed in cancer cells suggests unique cell specific mechanisms of regulation. Such an idea is further supported by our proteomics study wherein beta-arrestin 1 despite being a bonafide target of miRNA-7 is not altered in the cardiomyocytes [30] similar to EGFR indicating that cellular studies to assess miRNA targeting may not reflect the unique cardiomyocyte specific regulation. This suggests that a validated target in one cell system may not be viable target in another terminally differentiated cell suggesting an important and underappreciated role for cell-specific RNA binding proteins that could regulate the action of miRNAs. Together our study unravels an interesting aspect that despite multiple predicted targets for miRNA, only a few targets are significantly altered which may contribute to the overall phenotype.

## Acknowledgments

This work is supported by Postdoctoral Fellowship Grant, AHA, 10POST3610049 (MKG) and S10ODO21561 to SVNP. Furthermore, the proteomics core of Lerner Research Institute us supported by S10OD023436.

## Disclosure

The authors of the paper have no conflicts and nothing to disclose.

## Abbreviations

EGFR: Epidermal growth factor receptor
ERBB2: Erythroblastic oncogene
Erbb2: Proto-oncogene Neu
TAC: Transverse aortic constriction
DCM: Dilated Cardiomyopathy

## Declarations

### Funding

This work is in part, supported by Postdoctoral Fellowship Grant, AHA, 10POST3610049 (MKG). Conflicts of interest/Competing interests: The authors have not conflicts interest to disclose.

### Ethics approval

All the animal studies were approved by the Cleveland Clini IACUC.

### Availability of data and material

Primary data including global cardiac proteomics data is available and the authors will comply with reasonable requests.

### Authors’ contributions

Manveen K Gupta, Anita Sahu, Xi Wang, Elizabeth E. Martelli, Kate Stenson and Sathyamangla V. Naga Prasad - Designed, performed and optimized experiments, analyzed and interpreted data. Manveen K Gupta, Anita Sahu and Sathyamangla V. Naga Prasad - Drafted and edited the final manuscript..

## Supplementary Figure Legends

**Supplementary Figure 1.**
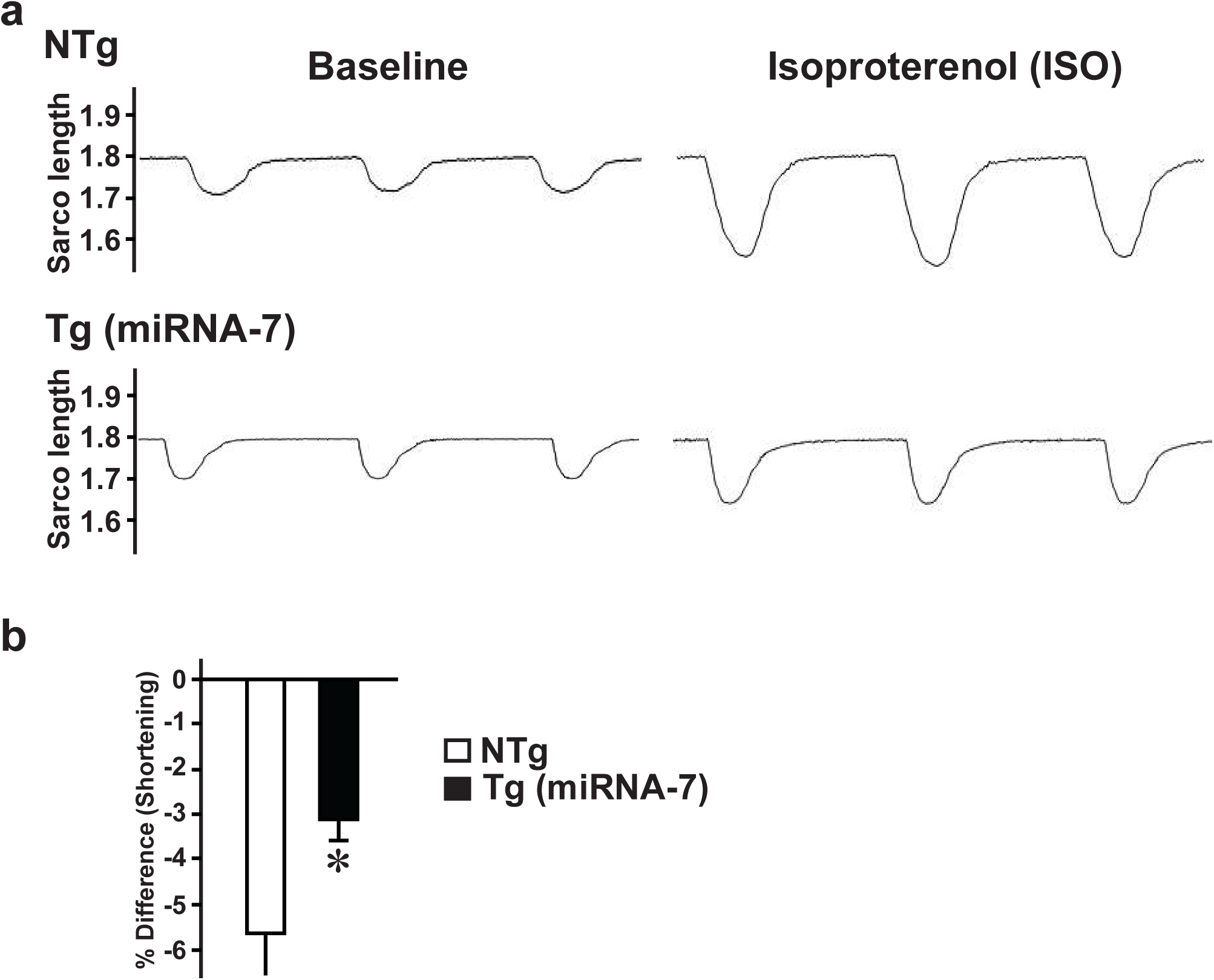
Adult cardiomyocytes were isolated from 3 months old NTg and Tg mice and were paced at 1 Hz (using the IonOptix Myopace system) to record the baseline cardiomyocyte contraction. Following steady baseline contractions, the cells were stimulated at 10 μM isoproterenol (ISO) and cardiomyocyte contractions were continously recorded using IonOptix System. **a**, Baseline myocyte contractility in NTg (upper left panel) and in vitro ISO-stimulated myocyte contractility (upper right panel). Baseline contractility in Tg (lower left panel) and in vitro ISO-stimulated myocyte contractility (lower right panel). **b**, % change in sarcomere length in response to in vitro ISO stimulation (n=8 mice each Tg or NTg; 15-20 adult cardiomyocytes from each NTg or TG mice).

**Supplementary Figure 2.**
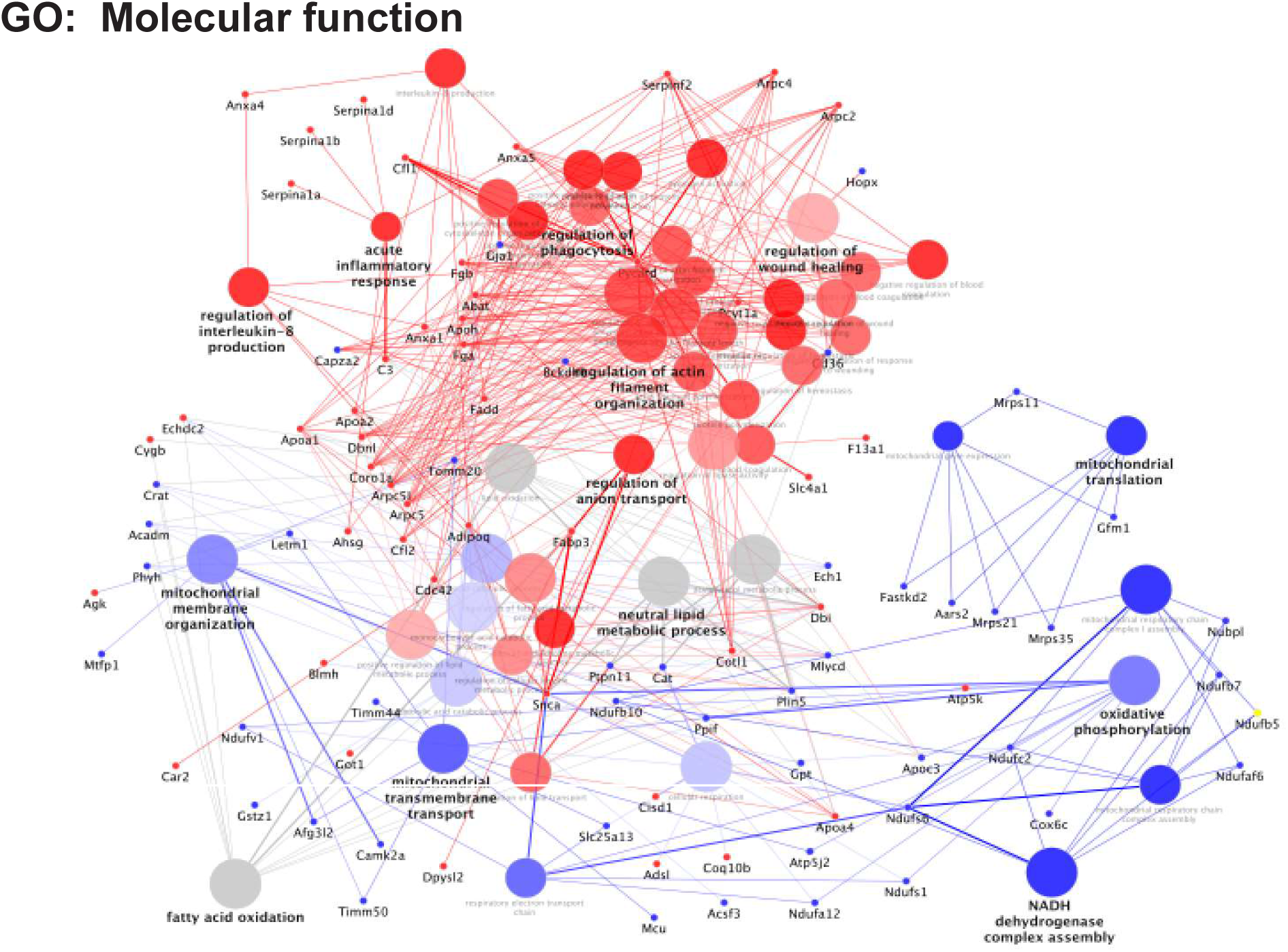
Gene ontology (GO) molecular functional networking analysis representing the higher magnification iteration of Figure 5b.

**Supplementary Figure 3.**
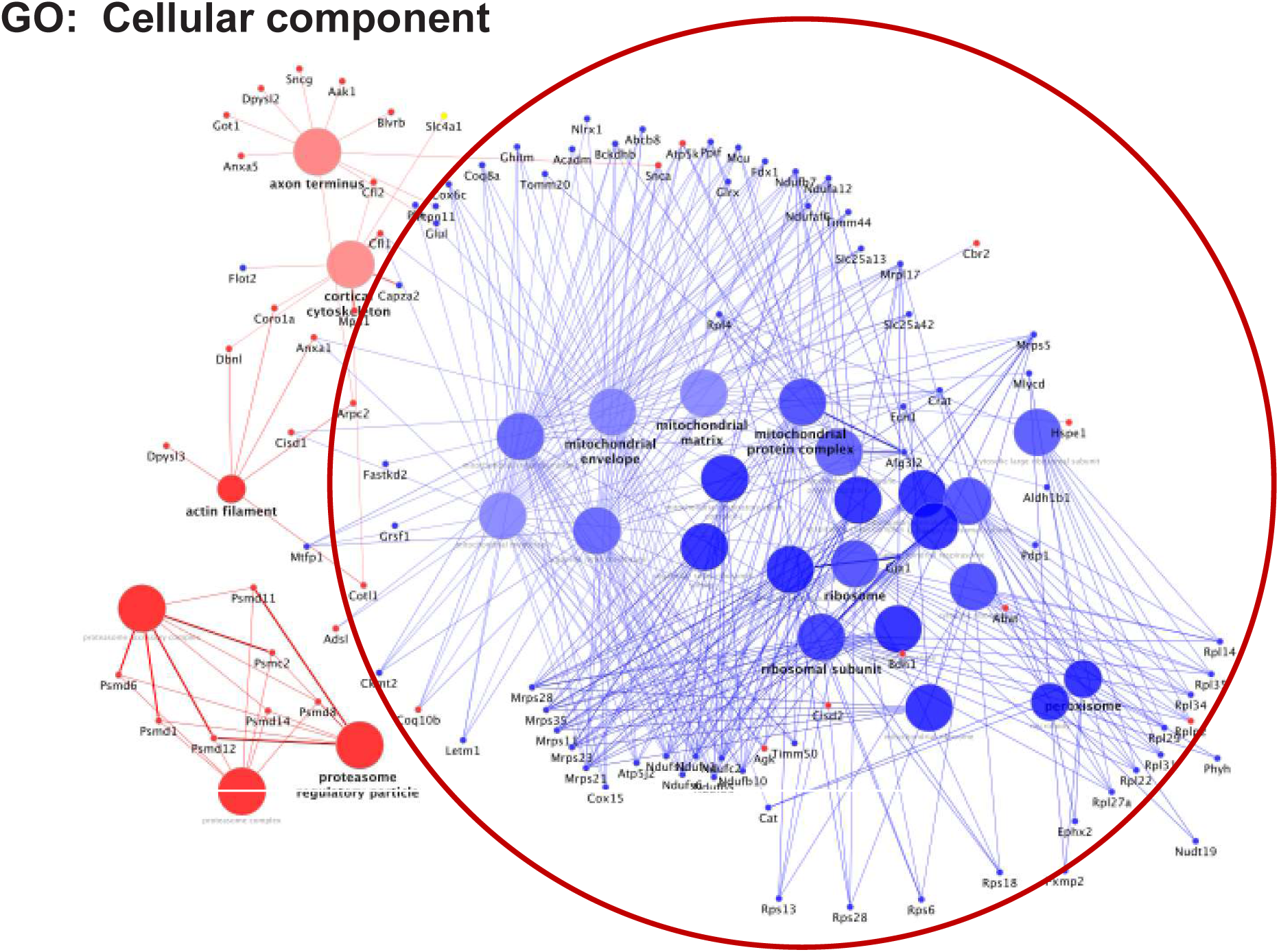
Gene ontology (GO) cellular component analysis representing the higher magnification iteration of Figure 5c.

## Notes

### Competing Interest Statement

The authors have declared no competing interest.

